# An optimized method for high-quality RNA extraction from distinctive intrinsic laryngeal muscles in the rat model

**DOI:** 10.1101/2022.03.23.485541

**Authors:** Angela M. Kemfack, Ignacio Hernández-Morato, Yalda Moayedi, Michael J. Pitman

## Abstract

Challenges related to high-quality RNA extraction from post-mortem tissue have limited RNA-sequencing (RNA-seq) application in certain skeletal muscle groups, including the intrinsic laryngeal muscles (ILMs). The present study identified critical factors contributing to substandard RNA extraction from the ILMs and established a suitable method that permitted high-throughput analysis. Here, standard techniques for tissue processing were adapted, and an effective means to control confounding effects during specimen preparation was determined. The experimental procedure consistently provided sufficient intact total RNA (*N* = 68) and RIN ranging between 7.0 and 8.6, which was unprecedented using standard RNA purification protocols. This study confirmed the reproducibility of the workflow through repeated trials at different postnatal time points and across the distinctive ILMs. High-throughput diagnostics from 90 RNA samples indicated no sequencing alignment scores below 70%, validating the extraction strategy. Significant differences between the standard and experimental conditions suggest circumvented challenges and broad applicability to other skeletal muscles. This investigation remains ongoing given the prospect of therapeutic insights to voice, swallowing, and airway disorders. The present methodology supports pioneering global transcriptome investigations in the larynx previously unfounded in literature.

## Introduction

Vocal fold (VF) motion is regulated by the intrinsic laryngeal muscles (ILMs), which facilitate glottic activities integral to daily life, including voicing, swallowing, and breathing^1^. To perform these roles, the ILMs control VF movement via complex neuromuscular activation^2, 3^. Upon contraction, the posterior cricoarytenoid (PCA) abducts the VFs, opening the glottis for swallowing and airflow during active and passive breathing^3^. In contrast, the thyroarytenoid (TA) muscles within the adductor complex adduct the VFs, positioning the vocal folds for phonation^4^. Each set of muscles is innervated by different branches of the recurrent laryngeal nerve (RLN). Following nerve injury, newly-formed axons from the proximal stump grow back to these muscles in an aberrant fashion, being directed in part by guidance cues. This non-selective reinnervation can result in synkinesis and VF paralysis with significant patient morbidity^4–9^. Effective treatment of nerve paralysis to re-establish normal VF function will require the restoration of the pre-injury innervation pattern. Further, therapeutic manipulation of the guidance cues expressed by the ILMs remains essential for rehabilitation. Evaluation of gene expression in these muscles before and after RLN injury will provide the insight necessary to identify high-value targets for further research and therapeutic intervention.

Pathological investigation of laryngeal muscle tension disorders is particularly favorable in the rat, a well-studied model of RLN injury and vocal fold paralysis^8^. Unfortunately, ILMs have not been amenable to high-throughput assessments as readily as other soft tissue types due to technical and biological challenges^10^. The impact of these challenges on the RNA extracted from skeletal muscles, including the ILMs, is consistent across species^11^. For instance, human post-mortem skeletal muscle analysis previously provided substandard RNA quality, summarized as RNA integrity numbers (RIN) ranging between 1.6 and 7.6. Given these results, only low-throughput analysis of the skeletal muscles was feasible^11^. Accordingly, suitable high-quality RNA extraction methods from the distinctive ILMs remain undocumented; hence, our limited understanding of the regulatory pathways that distinguish these muscles.

Identifying factors that hinder global transcriptome analysis of the rat ILMs was rarely supported in the literature before the present study. Moreover, gene expression assessment of the ILMs previously relied upon low-throughput profiling technologies (e.g., qRT-PCR, microarrays) and histological studies due to their lower requisites for RNA quality (McCullen 2009)^12–19^. Whole-transcriptome RNA-Seq provides a broader characterization and distribution of mRNA transcripts and their isoforms and thus requires highly undegraded RNA samples^20^.

The relevant challenges of high-quality RNA extraction are likely due to the small size, unique cellular content, and fiber composition of the rat ILMs^21^. Critical variables that influenced the quality of the RNA also included the conditions of the tissue, or the impact of degradation, during processing steps. The vulnerability of RNA in rat ILMs contradicts findings that characterize skeletal muscle mRNA as highly stable relative to the liver and brain^10, 11^. These evaluations suggested that variable RNA stability across species, tissue types, and organ layers may require modified extraction strategies according to the specimen and sample size. Previous studies have successfully performed a high-throughput analysis of the human^22^ and rabbit^23, 24^ VFs. Of relevance, these investigations did not perform an exhaustive analysis of the ILM tissue profiles. Rather than isolated ILM alone, they studied all the soft tissue associated with the VFs, including VF mucosa and respiratory epithelium. A recent Parkinson’s study demonstrated RNA-Seq expression analysis from the rat TA; however, their sequencing strategy enabled the input of low-throughput RNA extracts^25^. This method, commonly known as TruSeq, requires a small sample size and has unique RNA specifications unamenable to other bulk RNA-Seq technologies^26^. Despite these advancements, there remains a lack of publically available data describing a functional and reliable extraction method from the distinctive ILMs for general RNA-Seq use.

The present study aimed to establish a systematic, high-quality RNA extraction method from isolated postnatal rat ILMs that permitted RNA-Seq. Our lab developed a suitable technique to minimize native and non-native derived degradation and enhance RNA recovery. This study also identified an ideal weight model to limit the impact of total tissue weight during handling steps. Collectively, these findings demonstrate the critical factors of RNA isolation techniques that permit high-quality transcriptome analysis.

## Results

### Comparison of RNA extraction protocols

Standard extraction protocols were evaluated for the consistent production of high-quality RNA from isolated PCA, lateral thyroarytenoid (LTA), and medial thyroarytenoid (MTA) intrinsic laryngeal muscles. This study quantified the impact of mitigation strategies to establish optimal tissue processing conditions (a comprehensive workflow of the experimental design is summarized in **Fig. 1**). UV spectrophotometry (Nanodrop) was used to quantify RNA purity (A_260_/A_280_ ratio), concentration (ng/µL), and total RNA yield (ng); while electropherogram (bioanalyzer) analysis provided RIN values for quality control (QC) assessment. RNA extraction using the following kits were performed according to the manufacterer’s instructions, unless otherwise stated. In compliance with RNA-seq recommendations, this study aimed to generate extracts with greater than 500 ng Total RNA, 1.98 A_260_/A_280_ ratio, and 7.5 RIN. Qualifications for bioanalyzer assessment included sufficient total RNA recovery (> 200 ng), high purity (≥ 1.90), and no chemical contamination detection (**Table 1**).

**Table 1.** Assessment of protocols for RNA extraction from the rat intrinsic laryngeal muscles (ILMs). An analysis of variance (one-way ANOVA followed by Tukey’s post-hoc test) was used to compare the commercial protocols to the experimental method (*P* < 0.0001). The *pt* indicates the number of pooled tissues per extract, while *n* represents the number of replicates The RNA concentration (ng/μl) (shown as ng total RNA) and 260/280 ratio were measured using Nanodrop Spectrophotometer. Data are presented as mean ± SD; P < 0.05. RNA degradation was assessed using Agilent 2100 Bioanalyzer. The selection for Bioanalyzer indicates the quantity of samples applicable for RIN (RNA integrity number) analysis. Individual sample qualifications for QC were selected based on the starting material requirements for RNA-Seq (i.e., greater than 200 ng total RNA and 1.90 purity) and the presence of impurities in the extract. N/A RIN denotes samples RIN values that were not calculable.

|  |  | Samples<br>(n) | Total RNA (ng) | 260/280 ratio | Samples<br>for RIN | Positive<br>RIN | N/A<br>RIN | RIN value |
| --- | --- | --- | --- | --- | --- | --- | --- | --- |
| **Quick-RNA**<br>**MicroPrep** | (pt = 1) | 6 | 247.76 ± 107.72<br>(p = 0.94) | 2.05 ± 0.09<br>(p > 0.99) | 0 | - | - | - |
|  | (pt = 2) | 9 | 791.69 ± 294.06<br>(p > 0.99) | 1.63 ± 0.61<br>(p = 0.12) | 0 | - | - | - |
|  | **Total** | 15 | 574.11 ± 360.05<br>(p = 0.99) | 1.80 ± 0.51<br>(p = 0.32) | 0 | - | - | - |
| **RNeasy**<br>**Micro** | (pt = 1) | 6 | 94.07 ± 68.61<br>(p = 0.79) | 3.60 ± 2.91<br>****<br>(p < 0.0001) | 0 | - | - | - |
|  | (pt = 2) | 24 | 700.09 ± 529.43<br>(p > 0.99) | 2.12 ± 0.16<br>(p = 0.99) | 0 | - | - | - |
|  | **Total** | 30 | 578.89 ± 532.83<br>(p = 0.90) | 2.42 ± 1.36<br>(p = 0.24) | 0 | - | - | - |
| **TRIzol**<br>**Reagent** | (pt = 1) | 45 | 2458.84 ± 2070.48<br>****<br>(p < 0.0001) | 1.52 ± 0.10<br>****<br>(p < 0.0001) | 0 | - | - | - |
|  | (pt = 2) | 18 | 4148.15 ± 1996.87<br>****<br>(p = 0.0001) | 1.57 ± 0.09<br>**<br>(p = 0.002) | 0 | - | - | - |
|  | **Total** | 63 | 2941.50 ± 2174.27<br>****<br>(p < 0.0001) | 1.54 ± 0.1<br>****<br>(p < 0.0001) | 2 | 1 / 2 | 1 / 2 | 7.0 ± 0.0 |
| **½ TRIzol**<br>**Reagent** | (pt = 1) | 12 | 4430.11 ± 4049.31<br>****<br>(p = 0.0001) | 1.51 ± 0.11<br>**<br>(p = 0.004) | 0 | - | - | - |
| **TRIzol &**<br>**RNeasy**<br>**Micro** | (p t= 1) | 3 | 1309.35 ± 355.11<br>(p > 0.99) | 1.54 ± 0.04<br>(p = 0.59) | 0 | - | - | - |
| **TRIzol &**<br>**PureLink**<br>**RNA Mini** | (pt = 1) | 51 | 688.65 ± 611.70<br>(p = 0.97) | 1.67 ± 0.13<br>****<br>(p < 0.0001) | 19 | 12 / 19 | 7 / 19 | 4.2 ± 1.3<br>****<br>(p < 0.0001) |
| **Aurum** | (pt = 1) | 15 | 284.96 ± 165.61<br>(p = 0.58) | 2.42 ± 0.35<br>(p = 0.63) | 0 | - | - | - |
|  | (pt = 2) | 27 | 311.47 ± 260.37<br>(p = 0.29) | 2.25 ± 0.62<br>(p > 0.99) | 0 | - | - | - |
|  | **Total** | 42 | 302.01 ± 229.18<br>(p = 0.11) | 2.31 ± 0.54<br>(p = 0.78) | 15 | 15 / 15 | - | 5.2 ± 1.4<br>****<br>(p < 0.0001) |
| **Aurum &**<br>**RNAlater** | (pt = 3) | 48 | 658.80 ± 330.17<br>(p = 0.96) | 2.27 ± 0.42<br>(p = 0.97) | 25 | 25 / 25 | - | 8.0 ± 0.4<br>(p = 0.68) |
| **Proposed**<br>**method** | (pt = 3) | 68 | 981.28 ± 446.11 | 2.14 ± 0.09 | 68 | 68 / 68 | - | 7.8 ± 0.4 |
| (Aurum) |  |  |  |  |  |  |  |  |

**Table 2.** An overview of the specifications of the evaluated commercial kits and protocols

|  | **QuickRNA Microprep** | **RNeasy Micro** | **TRIzol Reagent** | **TRIzol + RNeasy Micro** | **TRIzol + PureLink RNA Mini** | **Aurum Total RNA Mini** |
| --- | --- | --- | --- | --- | --- | --- |
| Starting material | $\leq 10$ mg tissue (stored at $-80^{\circ}\text{C}$) | $< 5$ mg tissue (stored at $-80^{\circ}\text{C}$) | 50-100 mg tissue (stored at $-80^{\circ}\text{C}$) | $< 15$ mg tissue (stored at $-80^{\circ}\text{C}$) | $\leq 200$ mg tissue (stored at $-80^{\circ}\text{C}$) | $\leq 40$ mg thawed tissue (stored at $-80^{\circ}\text{C}$) |
| Approx. preparation time | $\sim 20$-30 min | $\sim 30$-40 min | $\sim 1$-2 hr | $\sim 1$-2 hr | $\sim 60$ min | $\sim 30$-60 min |
| Binding capacity | 10 $\mu\text{g}$ total RNA | 100 $\mu\text{g}$ total RNA | N/A | 100 $\mu\text{g}$ total RNA | 1000 $\mu\text{g}$ total RNA | $> 100$ $\mu\text{g}$ total RNA |
| Storage | RT | RT | 15-30$^{\circ}\text{C}$ | 15-30$^{\circ}\text{C}$/ RT | 15-30$^{\circ}\text{C}$/ RT | RT |
| DNase digestion | Yes | Yes | No | No | No | Yes |
| Elution volume | 20 $\mu\text{l}$ | 10-14 $\mu\text{l}$ | N/A | 20 $\mu\text{l}$ | 30 $\mu\text{l}$ | 30 $\mu\text{l}$-2 × 40 $\mu\text{l}$ |
| Advantages | Rapid procedure | Rapid procedure | High RNA yield | High RNA yield | Reduced contamination | Purity $> 1.96$ |
| Disadvantages | Inconsistent recovery<br>Low RNA yield | Inconsistent recovery<br>Low RNA yield | Low Purity<br>Phenol contamination | Low purity<br>Phenol contamination | Low purity<br>Phenol contamination | Purity $> 2.2$<br>Low RNA yield |

#### *Quick*-RNA™ MicroPrep

The *Quick*-RNA MicroPrep protocol (Zymo Research) provided a cost-effective means for performing RNA isolations. This column kit can be favorable for RNA extraction because it allows for polynucleotide isolation void of phenol contamination. Here, tissue samples were thawed and subsequently manually disrupted using RNase-free homogenization tubes. In preliminary evaluations of this protocol, favorable purities were observed (2.05 ± 0.09) (mean ± SD) using only one tissue sample *(pt=*1, where *pt* denotes pooled tissue); however, the amount of recovered RNA was low (247.76 ± 107.72 ng) and often did not meet the 200 ng requirement for RNA-Seq application (**Table 1**). To mitigate the low RNA yield, ILMs were pooled (*pt*=2), prompting both an increase in total RNA output (791.69 294.06 ng, *P* > 0.99) and a depreciation in the overall A_260/_ A_280_ ratio (1.63 ± 0.61, *P* = 0.99) that were not statistically significant. Additionally, substantial variations in the quality of samples across assays discouraged the continued use of this protocol (**Table 1**).

#### RNeasy^®^ Micro Kit

The RNeasy Micro Kit (Qiagen) was performed under identical conditions for tissue processing as previously stated. Advantages of the protocol included the absence of phenol detection in the recovered RNA and the short total procedure time. Using individual, frozen ILMs as starting material (*pt*=1) resulted in high variance in the A_260_/A_280_ ratio (3.60 ± 2.92) and amount of recovered RNA (94.07 ± 68.61 ng) such that extracts performed in the same batch were inconsistent and rarely qualified for further processing (**Table 1)**. Though increased tissue pooling (*pt*=2) resulted in a statistically significant difference in the overall A_260_/A_280_ ratio (*P* < 0.0001), the increase in total RNA yield (*P* > 0.99) was nonsignificant (**Table 1**). This kit was discarded due to considerable discrepancies in RNA quantity and purity across assays (**Table 1**).

#### TRIzol Reagent

Trizol reagent (Invitrogen) yielded significantly higher total RNA quantities using individual ILMs compared to the results achieved using the spin-column kits (2458.84 ± 2070.48 ng, *Quick*-RNA, *P* = 0.05; RNeasy, *P* = 0.004). Limitations of this protocol included low A_260_/A_280_ ratios (1.54 ± 0.10) in the recovered extracts, a 96.8% incidence (61 out of 63 samples) of phenol contamination across assays, and substantially longer procedure time than column-based methods; however, this procedure regularly provided RNA yields that exceeded the RNA-Seq sample requirements (**Table 1**). The cell lysis method was modified to mitigate the technical challenges. Optimal physical disruption parameters required rest periods flanking the manual lysing periods. To limit contamination, chloroform was reduced by 10-20 μL per 1 mL of Trizol in 16 out of 63 samples. This study hypothesized that this reduction would account for the loss of guanidinium thiocyanate solution (TRIzol) during rotor-stator disruption procedures. Here, differences in the overall purity (*P* > 0.99) remained nonsignificant; however, this method provided the only uncontaminated samples in this trial. In addition, a slight reduction in the ratio of chloroform and prolonged incubation in isopropanol (∼20 min) resulted in an individual RNA extract (1 out of 16 samples) with spectrophotometer measurements suitable for further processing (A_260_/A_280_ = 1.92; Total RNA = 7836.89 ng; phenol contamination = 0). Bioanalyzer assessment indicated a 7.0 RIN; however, these changes failed to provide comparable results beyond this sample.

#### ½ TRIzol Reagent

Due to the promising results from reducing the amount of chloroform, this study examined the effect of further minimization of reagent. Hence, the volume ratio was altered to reduce the level of tissue exposure to the phenol-containing chloroform. Scaling down to 0.5X volumes of the reagents failed to reduce the incidence of contamination (100% phenol detection) and significantly increase the overall purity (*P* > 0.99) relative to the former method (**Table 1**). As a result, this study terminated the continued implementation of this procedure.

#### TRIzol Reagent + RNeasy Micro Kit

Following the Furtado method^27^, the present study integrated the guanidinium thiocyanate solution (TRIzol) and spin-column RNA purification (RNeasy) protocols to assess the moderation of purity and contamination. This technique did not reduce the detection of impurities (100% phenol contamination) or significantly increase the A_260_/A_280_ ratios (*P* > 0.99) and thus further trials were discontinued (**Table 1**).

#### TRIzol Reagent + PureLink RNA Mini Kit

Combination of the guanidinium thiocyanate solution (TRIzol) and PureLink RNA Mini Kit (Invitrogen) protocols resulted in enhanced RNA quality. Phase separation tubes were utilized to sequester the aqueous phase^28^, reducing the incidence of phenol contamination by 47.3% compared to the guanidinium thiocyanate solution alone. (26 out of 51 samples). However, this modified protocol elicited a statistically significant decrease in RNA yield compared to the standard protocol (*pt*=1, *P* < 0.0001, **Table 1**). The moderations applied here also increased the overall A260/A280 ratios to 1.67 ± 0.13 (*P* = 0.99) though this change was not statistically significant. This method provided several samples with spectrophotometer measurements that warranted bioanalyzer assessment despite the lower total RNA yield (**Table 1**). Electropherogram analysis detected only one extract with a RIN > 7.0 (7.2 RIN), while most extracts contained relatively degraded RNA with RIN values ranging from 2.6 to 5.4. In addition, the quantity of non-calculable RIN values discouraged ongoing manipulation of this technique (**Table 1**). Consequently, the guanidinium thiocyanate solution was discarded because this study was unable to establish conditions to isolate RNA with acceptable purities using the reagent.

#### Aurum Total RNA Mini Kit

The Aurum kit (Bio-Rad) provided the most favorable A_260_/A_280_ ratios (2.31 ± 0.54) throughout this investigation (**Table 1**). Despite the nonsignificant differences in purity compared to other tested column-based methods (1.80 ± 0.51 for *Quick-*RNA MicroPrep, *P* = 0.72 vs. Aurum kit; 2.42 ± 1.36 for RNeasy Micro, *P* = 0.62 vs. Aurum kit) the resultant A_260_/A_280_ ratios using this spin-column (Aurum) kit did not extend below 1.96. Though most extracts were inapplicable for meaningful RNA-seq analysis, this kit provided no invalid RIN values (RIN = 5.19 ± 1.39, **Table 1**). These results indicated significantly increased RIN compared to the combined guanidinium thiocyanate (TRIzol + PureLink) method (*P* < 0.0001, **Table 1**). However, a shortcoming of this column kit was its feasibility to recover at least 500 ng of total RNA from a single ILM (*pt* = 1) or two pooled ILMs (*pt* = 2, **Table 1**). This protocol was selected for further optimization due to its more manageable complications comparatively. The present study hypothesized that modified specimen conditions and starting material (*pt*=3) would improve RNA integrity yield, respectively. While the applied changes significantly increased the RIN (*P* > 0.0001), these changes were insufficient to promote protocol systemization or consistent recovery (**Table 1**).

### Impact of RNA stabilizing mechanisms

Stabilization of endogenous RNA by ribonuclease (RNase) inhibiting reagents improved RNA integrity compared to the standard method (**Fig. 2A**). The applied RNase inhibiting reagent (RNA*later*), which hinders RNase activity through precipitation and metal chelation^10^, significantly increased the RIN in the overall model (*P* < 0.0001, **Fig. 2A, Table 1**). RNA isolated under standard conditions resulted in considerable RNA degradation shown by the migration of noisy 28S/18S ribosomal peaks (**Fig. 2B-D, *left*)**. Electropherogram peaks showed general depreciation of RNA molecules and insufficient total RNA recovery using standard tissue handling procedures (**Fig. 2B-D, *left***). Alternatively, the addition of RNase inhibiting reagents caused more distinct ribosomal RNA peaks and increased the RIN (**Fig. 2B-D, *right***). Low 28S/18S ratios (below 2.0) also revealed the vulnerability of RNA during isolation steps; however, the reliability of the 28S/18S ratios as an indication of relative degradation is unclear (**Fig. 2B, D**). Together, the low 28S/18S ratios and the small, rounded peaks at the baseline observed in the electrophoretic traces indicated that RNase inhibiting reagent treatment provided partial stabilization (**Fig. 2B-D, *right***). Beyond its indispensable inhibitory properties, overnight incubation in RNase inhibiting reagent also promoted the hardening of the muscle fibers, which increased the labor required to affect cell lysis.

### Homogenization parameters for high-quality RNA

Developing an effective cell lysis method for the adult rat ILMs required systematic assessment of manual and mechanical homogenization strategies. Manual techniques involved physical breakdown via disposable pestle in specialized RNase-free homogenization tubes.

Mechanical rotor-stator homogenization (TissueRuptor II) provided high-speed, automated tissue breakdown through replaceable probes. Finally, an integrated lysis method that incorporated both strategies was also adopted.

Evaluation of different homogenization methods suggested RNA yield and RIN were dependent on the cell lysis method, while purity was not (*P* = 0.33). Across global experiments (*pt* =3), the integrated homogenization method increased the recovery of RNA (887.59 ± 216.18 ng) and demonstrated significant differences compared to the manual (423.77 ± 143.98 ng, *P* = 0.001) and mechanical methods (372.15 ± 145.10 ng, *P* = 0.0004) (**Fig. 3A**). Physical homogenization provided nonsignificant but comparatively higher RIN (8.4 ± 0.1) than the combined (8.1 ± 0.1, *P* = 0.93) and the rotor-stator lysis methods (7.4 ± 0.6, *P* = 0.06) in the overall model (*P* = 0.59, **Fig. 3B**). Though not significant, the depreciation of RNA recovery and RIN at high rotor-stator speeds implicated the role of mechanical lysis in RNA degradation. The range of RNA quantity using this disruption technique (3.49-1535.24 ng) also supported findings that implicate aggressive rotor-stator use as a source of RNA damage^29^. Ultimately, the integrated lysis strategy at medium rotor-stator speeds provided the most favorable RNA extracts due to its propensity to promote considerable RNA recovery with less damage. This study advises manual homogenization if labor and limited RNA quantity are not a concern. Collectively, these findings demonstrated a critical dichotomy in the breakdown of ILMs, that is, resilient fibers and limited, fragile RNA.

### Establishing the ideal weight model

The evaluation of total RNA recovery attributable to the span of rotor-stator use (**Supp. Fig. 1**) suggested a critical range of breakdown that provides maximal RNA recovery. In our experimental trials, the assessment of optimal rotor-stator durations indicated variable periods of lysis between the ILMs, their anatomical location (i.e., right and left side), and the respective assay. The present study surmised that these discrepancies reflected the weight of starting material, a confounding variable in the cellular disruption process. Furthermore, investigators observed up to ten-fold differences in RNA quantity from -10 to 10 seconds changes in the rotor-stator span. An ideal weight model was developed by measuring total tissue weights (*pt* = 3) and evaluating the results from varying disruption times across several assays. These investigations allowed the present study to systematize the experimental procedure by identifying the upper and lower limits that ensured considerable intact RNA yields given the tissue characteristics. **Table 3** presents a range of weights and disruption times that provided the greatest, undiminished recovery. Similar impacts to the RIN were observed in some but not all cases; hence, this study only presents the total RNA yield, which revealed a more undeviating pattern.

**Table 3.** Recommended range of optimal disruption times using adult rat ILMs. The appropriate span of rotor-stator disruption attributed to the rodent stage, total weight of specimen and muscle type is shown. The listed ranges were collated based on extraction efficiency.

| **Muscle Type** |  |  |
| --- | --- | --- |
| **PCA** | **LTA** | **MTA** |

| *Mass (mg) | Time (s) | *Mass (mg) | *Mass (mg) | Time (s) | *Mass (mg) |
| --- | --- | --- | --- | --- | --- |
|  |  |  |  | 10 mg (n = 5) | 25 – 30 s |
|  |  | 20 mg (n = 2) | 35 s | 20 mg (n = 9) | 30 s |
|  |  | 30 mg (n = 4) | 35 s | 30 mg (n = 4) | 30 – 35 s |
| 40 mg (n = 9) | 65 – 70 s | 40 mg (n = 11) | 35 – 40 s | 40 mg (n = 2) | 30 – 35 s |
| 50 mg (n = 6) | 70 – 75 s |  |  |  |  |
| 60 mg (n = 3) | 75 s |  |  |  |  |

\*Total muscle weights using 3 pooled tissues.

Linear regression was implemented to determine the correlation between the ILM type, pooled tissue weight, period of rotor-stator lysis, and RNA yield (**Fig. 3C, F)**. This revealed opposing regressions between the PCA and TA muscles. More specifically, sufficient RNA recovery was only feasible at >1 min using the abductor muscle (**Fig. 3C, F**). Shorter rotor-stator durations did not provide lysates that could readily pass through the silica-membrane. In contrast, the total RNA in the TA muscles was reduced by prolong homogenization with optimal times ranging from 25 to 40 s. Diverging from the PCA muscle, raw data from the LTA and MTA muscles were compounded because it was observed that they required similar disruption times (R^2^ = 0.24, **Fig. 3D**). Given the larger size of the PCA muscles, their homogenization needed to be longer to break down the tissue and provide sufficient nucleic acid yields (R^2^ = 0.31, **Fig. 3C**). Visualization of the observed and theoretical quantiles indicated a normal distribution of the presented data (**Fig 3D, G**). The residual plots confirmed the accuracy of the linear regression (**Fig 3E, H**). The results suggest variation in RNA recovery was attributable to the tissue density, weight, and disruption span (**Fig. 3C-G**).

### Effects of cryogenic grinding on RNA quality

A simple instrument, which circumvented cross-contamination and thawing, was developed to facilitate the breakdown of tissue hardened by RNase inhibiting reagent (**Fig. 4**). Initial tissue grinding in liquid nitrogen maximized the capacity of the silica-membrane (40 ± 20 mg). In addition, it reduced the instance of column clogging and hole formation from insufficient homogenization. The adapted method for cryogenic disruption significantly improved the RNA recovery in the overall model (**Fig. 5A**). Generally, this modification raised RNA quantities to 1003.64 161.181 ng relative to the standard method (661.44 ± 73.31 ng, *P* < 0.0001). A significant increase in RNA yield was observed in the PCA (*P* = 0.03) and MTA (*P* = 0.003) muscles, while the changes observed in the LTA were not significant (*P* = 0.15, **Fig. 5A**). On the other hand, significant differences in RNA purity were revealed in the LTA compared to other ILMs (*P* = 0.004) and the overall model (*P* = 0.01, **Fig. 5B**). Despite observing slight reductions in the experimental RIN compared to standard conditions, two-way ANOVA analysis revealed no significant changes overall in RNA stability due to the applied method (*P* = 0.06, **Fig. 5C**). Significant differences attributable to cryogenic disruption were only demonstrated in the RIN from MTA muscles (*P* = 0.02, **Fig. 5C**). The MTA was the only muscle with no significant improvements to the 28S/18S ratios (*P* > 0.99, **Fig. 5D**). However, two-way ANOVA analysis indicated significantly increased 28S/18S ratios from cryogenic disruption across the overall muscle types (*P* = 0.004, **Fig. 5D**).

### Validating the applied tissue pooling strategy

Various pooling strategies (*pt* = 1-3) were evaluated to determine the optimal number of muscles per sample to isolate sufficient quantities of RNA for sequencing (**Fig. 6**). In initial trials using the preferred *Aurum Total RNA Mini Kit*, pooling one or two ILMs resulted in inadequate total RNA yields (**Table 1**). However, the progressive increase in RNA recovery in the aforementioned experimental steps reconsidered the need for tissue pooling, as such different conditions were tested. Two-way ANOVA analysis indicated significant differences in the total RNA yield attributable to tissue pooling (*P* < 0.0001, **Fig. 6A**). These evaluations suggested that consistent high RNA recovery (>200 ng) is not achievable using individual ILMs. Pooling few muscles, however, significantly enhanced RNA recovery (*pt=*1 vs 3, *P* = 0.0004, **Fig. 6A)**. Subsequent trials with two pooled ILMs (*pt* = 2) resulted in sufficient RNA quantities in most (88.9%) but not all cases (*P* = 0.06, **Fig. 6B)**. In contrast, pooling ILMs from three adult rats (300g) consistently produced the recommended concentrations for RNA-seq (100% of samples, N=18**, Fig. 6B)**. Trials with three muscles demonstrated sufficient extraction across the PCA (846.54 ± 336.87 ng), LTA (694.89 .1 ± 394.07 ng, *P* = 0.05), and MTA muscle (1121.35 ± 555.19 ng), which was not readily observed using fewer pooled muscles (**Fig. 6**). Moreover, the advantage of pooling three muscles over two is assurance the samples will qualify for downstream processing.

### An overview of the experimental protocol

The collective experimental techniques provided (*N* = 68) RIN values from 7.0 to 8.6 (7.8 ± 0.4) with no RNA yields below 200 ng (981.28 ± 446.11, **Table 1**). Ultimately, the applied experimental modifications in conjunction with the Aurum Total RNA Mini kit significantly improved RNA recovery compared to other evaluated protocols (**Table 1**). Functional RNA extraction from rat ILMs required RNase inhibiting reagents during surgical steps and overnight incubation for nucleic acid stabilization. High-quality RNA extracts were achieved through implementing the integrated (manual and mechanical) cell lysis strategy, establishing the ideal weight model, and adapting cryogenic disruption. Finally, pooling three ILMs permitted consistent and sufficient total RNA yields. The reproducibility of the experimental method was confirmed using P15 and P32 (∼115 g) rats, which validated the effectiveness of the experimental conditions (**Supp. Fig. 2**). **Table 4** summarizes RNA-Seq (NovaSeq) data analysis taken from 90 RNA samples, with RIN ranging between 7.0 to 9.3.

**Table 4.**
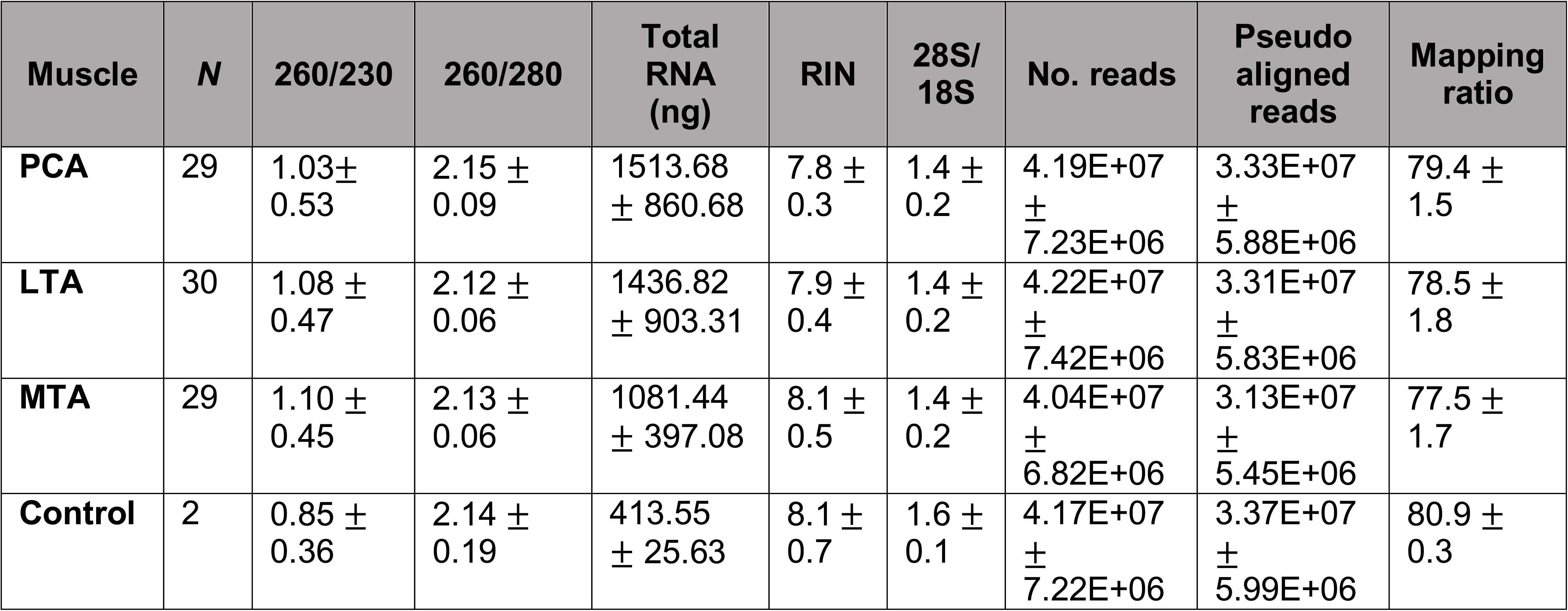
Summary of the downstream diagnostics from experimental total RNA extracts (*N* = 90). RNA samples were sequenced using a standard RNA-Seq pipeline (Illumina NovaSeq 6000), and sequencing reads were quantified using Kallisto. Note, the controls in this experiment were (1) ERNA extracts from the right ILMs aggregated and (2) RNA extracts from all paired ILMs combined .

| Muscle | N | 260/230 | 260/280 | Total RNA (ng) | RIN | 28S/18S | No. reads | Pseudo aligned reads | Mapping ratio |
| --- | --- | --- | --- | --- | --- | --- | --- | --- | --- |
| PCA | 29 | 1.03 ± 0.53 | 2.15 ± 0.09 | 1513.68 ± 860.68 | 7.8 ± 0.3 | 1.4 ± 0.2 | 4.19E+07 ± 7.23E+06 | 3.33E+07 ± 5.88E+06 | 79.4 ± 1.5 |
| LTA | 30 | 1.08 ± 0.47 | 2.12 ± 0.06 | 1436.82 ± 903.31 | 7.9 ± 0.4 | 1.4 ± 0.2 | 4.22E+07 ± 7.42E+06 | 3.31E+07 ± 5.83E+06 | 78.5 ± 1.8 |
| MTA | 29 | 1.10 ± 0.45 | 2.13 ± 0.06 | 1081.44 ± 397.08 | 8.1 ± 0.5 | 1.4 ± 0.2 | 4.04E+07 ± 6.82E+06 | 3.13E+07 ± 5.45E+06 | 77.5 ± 1.7 |
| Control | 2 | 0.85 ± 0.36 | 2.14 ± 0.19 | 413.55 ± 25.63 | 8.1 ± 0.7 | 1.6 ± 0.1 | 4.17E+07 ± 7.22E+06 | 3.37E+07 ± 5.99E+06 | 80.9 ± 0.3 |

## Discussion

The present study mitigated and quantified the factors that decreased total RNA recovery from rat vocal fold muscles. In identifying these significant contributors, this investigation established a systematic protocol that consistently produces high-quality RNA extracts. Our findings demonstrate that high-quality RNA extraction from isolated PCA, LTA, and MTA muscles is contingent on adapted tissue preparation steps. This research has pioneered a reproducible protocol that provides RNA extracts suitable for varied bulk RNA-Seq methods.

Comparisons of the purification procedures suggested that standard tissue preparation techniques, as per the respective protocol guidelines, do not account for variability or sensitivity of starting material. Moreover, the shortcomings of the tested spin-column kits were likely related to the assumed standardized preparation of animal tissue rather than a reflection of the effectiveness of the column-based systems. The low A_260_/A_280_ ratios and the inflated RNA yields observed with guanidinium thiocyanate solution (Trizol) indicated substandard extraction for high-throughput sequencing technology. Likewise, other laryngeal muscle studies that have used this reagent to extract RNA from the rat model have only performed low-throughput analysis^13–15, 30–32^. In selecting an appropriate kit, this study valued a ∼2.0 A_260_/A_280_ ratio and a low variation between samples in consideration of the influence of batch effect during RNA-Seq analyses^33^. The chosen kit provided the most favorable extracts with A_260_/_280_ ratios higher than 1.96, indicating successful DNase digestion. Consistent data void of chemical contamination across all assays also encouraged the use of this protocol. Though the standard method did not result in high RNA yields, it was the most practical complication to mitigate relative to other tested kits.

A limiting factor of the preferred kit was its inability to consistently provide sufficient total RNA without pooling muscle fibers (*pt* = 3). Compelling evidence in other rat model studies has demonstrated the feasibility of successful RNA-Seq analysis using the Aurum Total RNA Fatty and Fibrous Tissue Kit^25, 34^. Due to the greater column capacity (100 mg) of the aforementioned kit, its application can be assessed in future studies.

Assessment of recovered RNA revealed a significant influence of the tissue preparation procedures, where RNA degradation remained inevitable upon organismal death until the development of the lysate. In downstream assessments, degradation, depicted by the loss of short genes and the changes to longer genes in the mRNA transcript, perturbs the innate gene profile^35^. Similar deficiencies can also persist during the large-scale amplification of extracts with low RNA quantities. Previous studies have also evinced organismal death and the *post-mortem* interval (PMI) as external factors that mediate alterations to tissue transcriptomes^36–38^. Conversely, previous research has challenged the alterations to RNA integrity following prolonged ischemia finding no correlation between RNA quality and the PMI in human skeletal muscle^11^ and colonic epithelial tissue^39^. Without a clear consensus in the literature, the present study mitigated degradation during the PMI by minimizing the total procedure time and the use of RNase inhibiting reagents^38^. This study applied RNase inhibiting reagent (RNA*later*) to the dissected ILMs to limit degradation. Overnight incubation of the muscles in cold RNase inhibiting reagent established the immobilization of active hydrolytic contributors during freeze/thaw cycles. ILM dissection from fresh tissue rather than after incubation in the stabilizing reagent allowed for efficient separation. Storing individual ILMs also increased the surface area exposed to the RNase inhibiting reagent. Alternatively, incubating the whole larynx overnight would likely result in partial or fragmented muscle removal. Given the elevated RIN values and low 28S/18S ratios, this study concluded that the stability this reagent provided was not absolute. Previous methodological studies that assessed the efficiency of RNase inhibiting reagents at different time points demonstrated reduced RIN after 24 hours, justifying an overnight incubation^29, 40^.

Limitations of evaluations using RNase inhibiting reagent in this study were related to its effectiveness over time. Namely, overnight storage in RNase inhibiting reagents followed by prolonged storage at -80 °C was not assessed for changes in RNA integrity. Overnight storage was complicated by its propensity to cause column clogging and hole formation due to hardened ILMs that deemed several samples (23 out of 48) unusable (**Table 1**). Finally, due to the unreliability of the 28S/18S ratios, as shown in MTA muscles, this study could not draw any reliable conclusions apropos the efficiency of the reagent to enhance RNA stability. This study inferred RNA stability through visualization of the electropherogram peaks as such. Collectively, these results confirmed that this modification alone was insufficient for the consistent recovery of highly suitable RNA extracts.

The disruption paradigm developed here ameliorated the structural and functional challenges associated with high-quality RNA isolation. Establishing an optimal method that leads to the destruction of the cell membrane without considerable deterioration of intracellular components was a rate-limiting step throughout this study. Heat generation resulting from intensive rotor-stator disruption cycles contributed to increased degradation in our samples—given the linear relationship between increasing temperature and RNA denaturation^41^. The advantages of the developed method were four-fold. First, using an integrated homogenization strategy significantly increased the yield of RNA recovery per assay, allowing most extracts to qualify for RNA-Seq. Second, the ideal weight model permitted a systematic approach for optimal RNA extraction void of excessive inter-and intra-variability across the global experiments. Third, the conceptualization of the unique cryogenic disruption module provided an effective means for tissue disruption void of chemical and biological contaminants. Fourth, the experimental method reduced the incidence of column clogging and hole formation, making all recovered RNA applicable for bioanalyzer assessment (**Table 1**).

Limitations of the ideal weight model included observing no remarkable interaction between the span of rotor-stator disruption and the resultant RIN. Thus, because an accurate means to quantify the capacity of homogenization per ILM is unprecedented, the present method cannot guarantee extracts with RIN exclusively above 8.0. Alternatively, this protocol provides sufficient quantities of highly pure total RNA and RIN between 7.4 and 8.6 with limited variability compared to other skeletal muscles^11, 42^. The pronounced ILM sensitivity and the significant improvement of technical and biological challenges suggest the applicability of the experimental techniques to other skeletal muscles.

Although manual homogenization circumvented degradation from rotor-stator use, this technique alone provides insufficient RNA recovery for bulk RNA-Seq methods. Low RNA recovery can be problematic as it may lead to overamplification of sequences not representative of the entire transcriptome^43–45^. Manual homogenization is also more challenging to standardize due to the influence of human error. Maintaining consistent technical and biological conditions is essential for detecting outliers and variability of the sequencing data^22^. Given that increased variability may lead to unreliable RNA-Seq assessments, the present study recommends diligent manual lysis when a more sensitive library kit, such as TruSeq, is applied^46^. Otherwise, the integrated method and a reasonable approach to determine the ideal load for cellular breakdown provide maximization of RNA with sufficient RIN values needed for high-throughput analysis (**Table 1**).

Given the restrictions of the spin-columns (approximately 40 mg of starting material per column), pooling three ILMs per sample reflects both the minimum and the maximum amount needed for optimal RNA extraction of adult rat ILMs 100% of the time. Suitable RNA recovery is achievable using two ILMs, but only 88% of the time. A limitation to the proposed pooling strategy is the results may not accurately reflect population variation in gene expression levels^47^. Pooling studies that evaluated differential gene expression previously reported high false positivity rates and pooling bias from pooling less than 8 tissues^47^. Validation of RNA-Seq data using TruSeq, which notably allows for sequencing of degraded samples or low amounts of RNA, may be necessary to assess the impact of the experimental method^46^.

The challenges regarding effective extraction for RNA-Seq analysis reflected the profound influence of degradation and structural differences between the ILMs. The present study observed differences in susceptibility to RNA degradation relative to the specific ILM (**Supp. Fig. 1**). One hypothesis to consider was that the ILM structure impacts optimal processing conditions. Recent findings demonstrating increased resistance of the rat PCA and TA muscles to functional impairment and age-related changes supported this hypothesis due to their physiological roles^48^. This age-related study indicated that more resilient fibers in the PCA and TA muscles, relative to the cricoarytenoid muscle, are attributable to their inherent function. These observations suggested that the load required for the breakdown of the PCA is greater than that of the TA muscles due to their increased adaptive resistance and fiber content. Evaluating various purification strategies implicated degradation as particularly adverse for ILMs due to their small muscle fiber cross-sectional area and volume, ergo limited RNA abundance^49, 50^. Previous quantitative PCR analysis of abductor (PCA) and adductor (LTA and MTA) muscles also demonstrated the significant vulnerability of the cellular components of these muscles to damage during RNA isolation procedures^11^. In short, for optimal RNA extraction from rat ILMs, substantial force is needed to affect cell lysis; however, too much can compromise intact RNA (**Supp. Fig. 1)**. Finally, our findings suggested that the intrinsic characteristics of each muscle significantly impacted extraction. Thus a systematic approach must be used to identify the force that will optimize RNA extraction for individual ILMs.

### Conclusions

The present work demonstrates the hallmarks of critical factors that allowed for high-quality transcriptome analysis: stabilization of RNA provided by RNase inhibiting agents, integrated homogenization, the ideal weight model, adapted cryogenic disruption, and tissue pooling. High-quality RNA extraction from the ILMs for general RNA-Seq analysis required additional consideration of the low amounts of starting material, handling of the specimen, tissue specific composition, and tissue-specific degradation rates following death and excision. Comprehensive RNA-Seq analysis validated the efficacy of the experimental method identified herein, in general and specifically for rat vocal ILM..

## Methods

### Animals

This study was performed in compliance with the Public Health Service Policy on Humane Care and Use of Laboratory Animals, the National Institutes of Health Guide for the Care and Use of Laboratory Animals, and the Animal Welfare Act (7 U.S.C. et seq.). The Institutional Animal Care and Use Committee of Columbia University Medical Center approved the animal use protocol. Under a 12 h dark/light cycle and ambient temperature (21 °C), Sprague-Dawley rats were placed in group housing with *ad libitum* access to food and water. Young adult rats (∼300 g) were distributed in sets of triplicate treatment groups according to their sex (3 rats per group). The procedures and their modifications for the optimized protocol are described as follows:

### Tissue Preparation

Adult rats were euthanized by a lethal injection of ketamine and xylazine before laryngeal muscle dissection. Once euthanized, 70% ethanol served as a skin disinfectant before harvesting the larynx. A midline incision was performed in the neck. Following retraction of submandibular glands and extralaryngeal muscles, the larynx was identified and removed en bloc with the proximal trachea, esophagus, and pharynx^33^. The specimen was then were transferred to a sterile petri dish and rinsed in 2 mL of RNA*later*™ (catalogue #AM7021, Invitrogen, Vilnius, Lithuania), which served as a stabilizing agent. After aspirating excess solution from the petri dish, the specimen was placed onto a sterile surface for dissection via a Leica S8 APO compound microscope (Leica Microsystems, Buffalo Grove, IL). The esophagus and pharynx were carefully excised from the posterior aspect of the harvested tissue before extracting the right and the left laryngeal abductor muscles (PCA). The aryepiglottic folds were then removed to provide access for dissection of the right and left adductor muscles (LTA and MTA) with fine-tip forceps. ILMs from the rat larynx were less than 5 mm in length, and individual muscle bellies weighed between 2 mg and 13 mg. Rather than storing the whole larynx in the reagent after rinsing, per other optimized protocols for RNA purification, individual ILMs were immediately incubated in 1 mL of RNA*later* following dissection. Samples were then placed at 4°C overnight and then processed the following day.

### Assessment of RNA extraction protocols

The following kits were tested for the evaluation of the most effective RNA isolation: ZR *Quick-*RNA™ MicroPrep (catalogue #R1051, Zymo Research, CA,.), RNeasy^®^ Micro Kit (catalogue #74004, Qiagen, Hilden, Germany), TRIzol™ Reagent (catalogue #15596-026, Invitrogen Life Technologies, Carlsbad, CA, U.S.A.), and the Aurum™ Total RNA Mini Kit (catalogue #732-6820, Bio-Rad, Hercules, CA, U.S.A.). In addition, the following procedures were evaluated: TRIzol protocol using a half volume of all reagents, Trizol in combination with RNeasy Mini kit (Qiagen), Trizol plus PureLink™ RNA Mini Kit (catalogue #12183018A, Invitrogen, Carlsbad, CA, U.S.A.), and RNA*later* in sequential integration with the Aurum kit (Bio-Rad). RNA purification was followed according to the kit instructions, and deviations from the recommended preparation of the starting material are reported in the Results section. Protocol modifications to the Aurum kit and Trizol protocols facilitated the optimization of RNA purification. Here, parameters related to homogenization duration, disruption method, number of tissues per tube, and volume of reagents were modified

### Specimen Preparation

Before next-day isolation procedures, the benchtop was sprayed with RNase*Zap*™ (catalogue #AM9780, AM9782, Invitrogen, Waltham, MA, U.S.A.) for removing ribonuclease contamination. Other surfaces and instruments were also sprayed during all laboratory processes. The Aurum Total RNA Mini kit allowed for the most optimal and consistent extraction of high-quality RNA. For the prescribed procedure, tissues were removed from stabilizing reagent and briefly dabbed on a paper towel to remove excess solution, according to the manufacturer’s guidelines. Samples were pooled (*pt* = 3) according to muscle type, position (right or left), and gender. Similar laryngeal muscle tissues were combined in respective BioMasher III tubes (Nippi, Inc., Japan), with the filters removed, to facilitate manual grinding and lyophilization in liquid nitrogen via the disposable pestles. Before the physical breakdown of muscle tissue, the pooled samples were weighed to determine the appropriate duration of mechanical homogenization to yield cell lysates with significant intact RNA. After pooling, the calculated median total weights were 50 mg, 40 mg, and 20 mg for PCA, LTA, and MTA, respectively. An apparatus for cryogenic disruption was assembled using medium binder clips and BioMasherIII tubes (**Fig. 4**). After placing liquid nitrogen in the surgical tray, the pooled muscle fibers were broken down via disposable pestle. For further muscle dissociation, the RNA-preserving denaturant (lysis buffer) was added to the lyophilized powder where samples were subject to additional disruption through repeated pipetting, following the Aurum kit protocol. The processing of outlier values with robust weights (PCA) was ameliorated by passing the lysate through two columns and aggregating the extracts in the last step. The breakdown of any remaining particulates via the pestle facilitated blending of the lysate. Subsequently, mechanical homogenization ensued through rotor-stator disruption. Disposable probes were subject to cleansing in consecutive rounds of RNAse-free water and 70% ethanol before automated lysing to avoid cross-contamination between samples. In all samples disruption occurred in 1 min intervals to avoid heat generation derived from the TissueRuptor II (catalogue #9002755, Qiagen, MD)^29^.

### RNA Extraction

RNA concentration was measured using a micro-volume spectrophotometer NanoDrop ONE (catalogue #ND-ONE-W, Thermofisher, Wilmington, DE). RNA integrity, the value of which serves as the final determinator of efficacy for RNA-Seq analysis, was assessed using an Agilent 2100 bioanalyzer. The RNA integrity number (RIN) calculated using this software presents a range of values from 1.0 to 10.0—indicating complete degradation and intact RNA, respectively^51^. This study aimed to reach RIN close to or higher than 8 for meaningful genomic analyses. The optimized protocol systematically yielded at least 500 ng total RNA, approximately 200 ng is generally required for RNA-seq.

### Statistics

The present study expresses all data as mean ± SD unless otherwise stated. All statistical analyses were performed using GraphPad Prism 9.3.1 (San Diego, CA). Ordinary one-way ANOVA followed by Dunnett’s multiple comparisons tests were performed to compare the experimental results to the recovery of RNA using currently available protocols. A repeated analysis of variance followed by Tukey’s post-hoc tests allowed for individual comparisons of the commerical extraction methods. To examine the effect of RNA*later* and cryogenic disruption on RNA quality, two-way ANOVA followed by Bonferroni’s multiple comparisons test was computed. The assessment of homogenization techniques and pooling strategies were facilitated by standard two-way ANOVA followed by Dunnett’s multiple comparisons tests to assess RNA quantity and purity that resulted from cryogenic disruption. Linear regression analysis was performed to demonstrate the relationship of contributing factors in the ideal weight model. The data were tested for normality via Q-Q plot and regression analysis. Finally, data in all supplemental figures were computed by two-way ANOVA followed by Tukey’s post-hoc tests. The present study considered *P* < 0.05 as the level of statistical significance. (\**P* < 0.05; ** *P* < 0.01; *** *P* < 0.001; **** *P* < 0.0001)

### Data availability

The data used to substantiate all findings in this manuscript will be provided upon request.

## Acknowledgments

We thank Almudena Bosch, Medini Annavajhala, and Larissa Williams for contributing their expertise in RNA-related studies; and the JP Sulzberger Columbia Genome Center for their services and assistance in next-generation sequencing. This work was supported by grant 1R01DC018060-1 of the US. National Institutes of Health b to MJP.

## Contributions

### Conceptualization

A.M.K., I.H.M, M.J.P Methodology: A.M.K., I.H.M, Y.M., M.J.P. Formal analysis: A.M.K., I.H.M. Investigation: A.M.K. Resources: M.P. Data Curation: A.M.K Writing – Original Draft: A.M.K. Writing – Review & Editing: I.H.M, Y.M., M.J.P Visualization: A.M.K., Supervision: M.J.P Validation: A.M.K. Project Administration: M.J.P. Funding Acquisition: M.J.P

## Ethics declaration

Competing interests

The authors declare no competing interests.

## Additional Information

Publisher’s note

Publisher's note
|  |  |  |
| --- | --- | --- |
| **Kits** | **Purification of RNA** | **Selection for Bioanalyzer** |

**Supplementary Information**

**Rights and permissions**

**About this article**

**Comments**

---

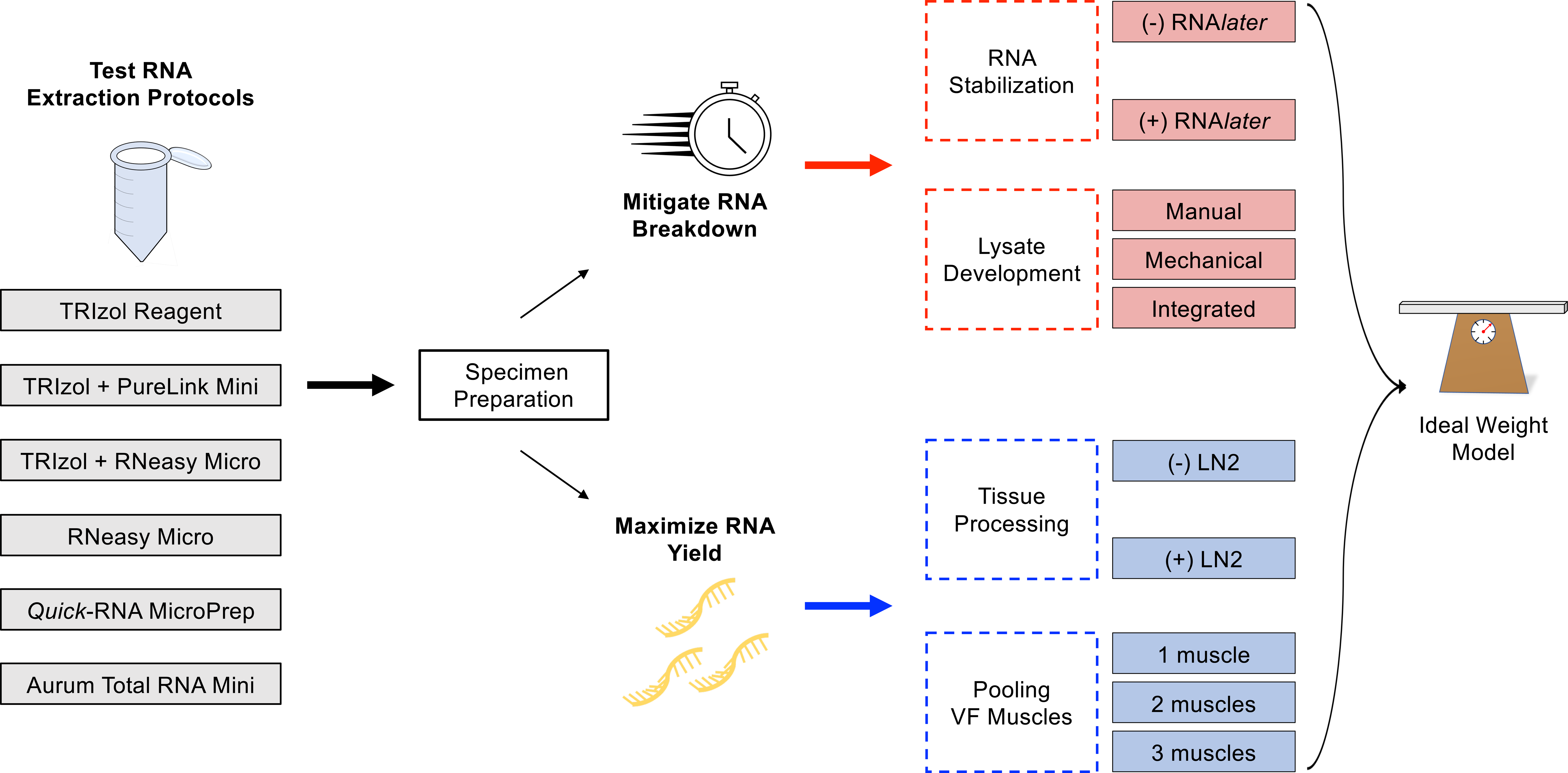

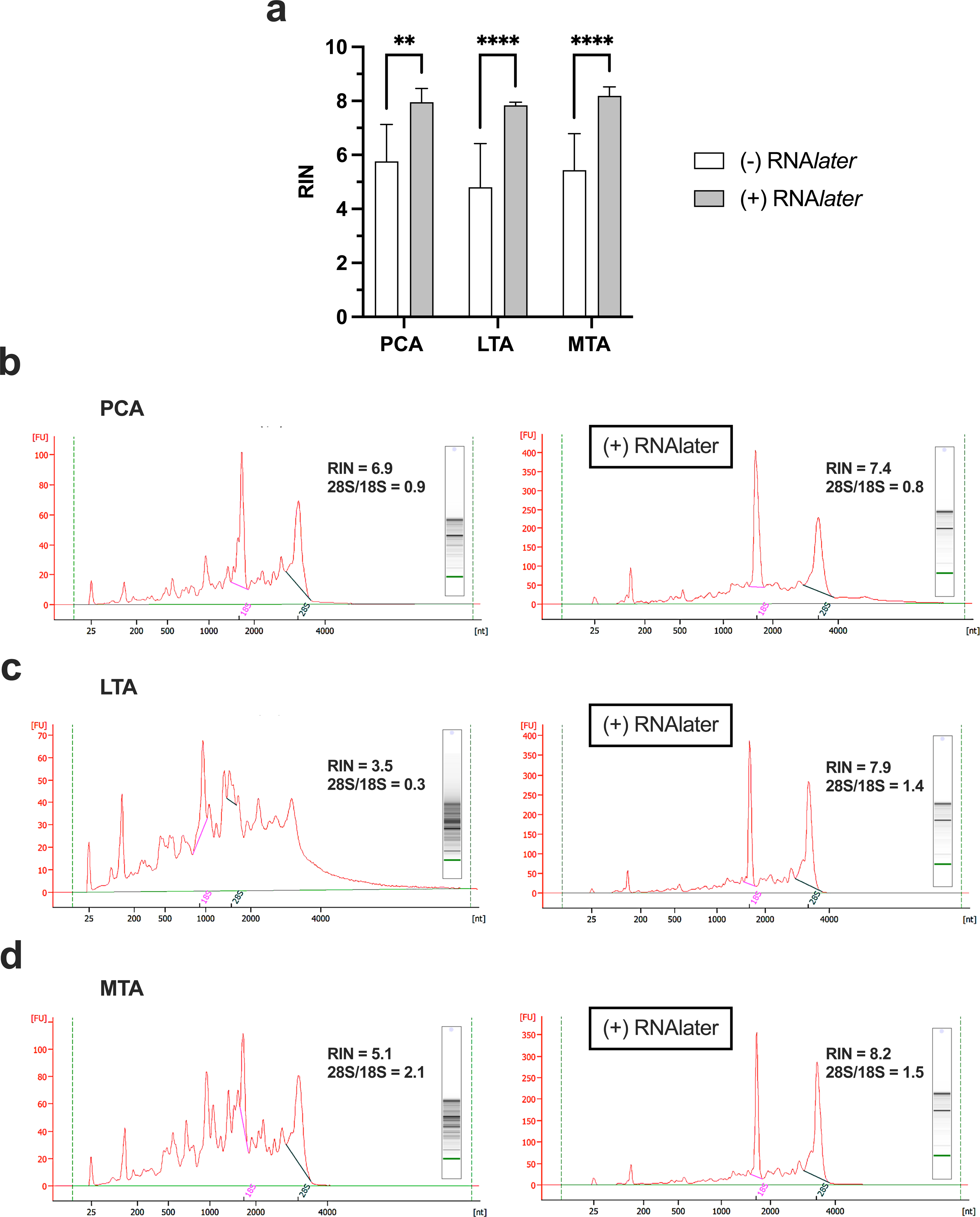

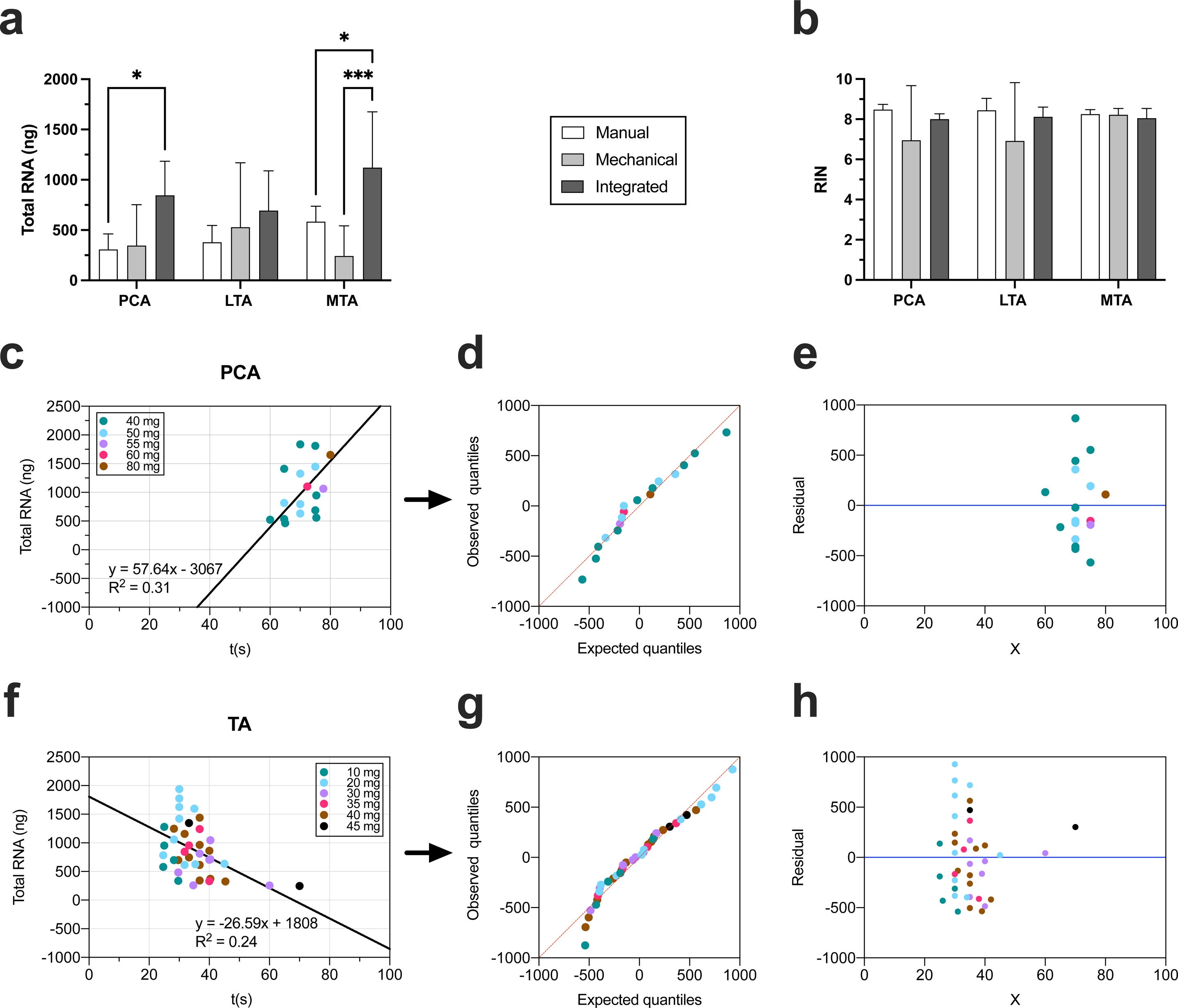

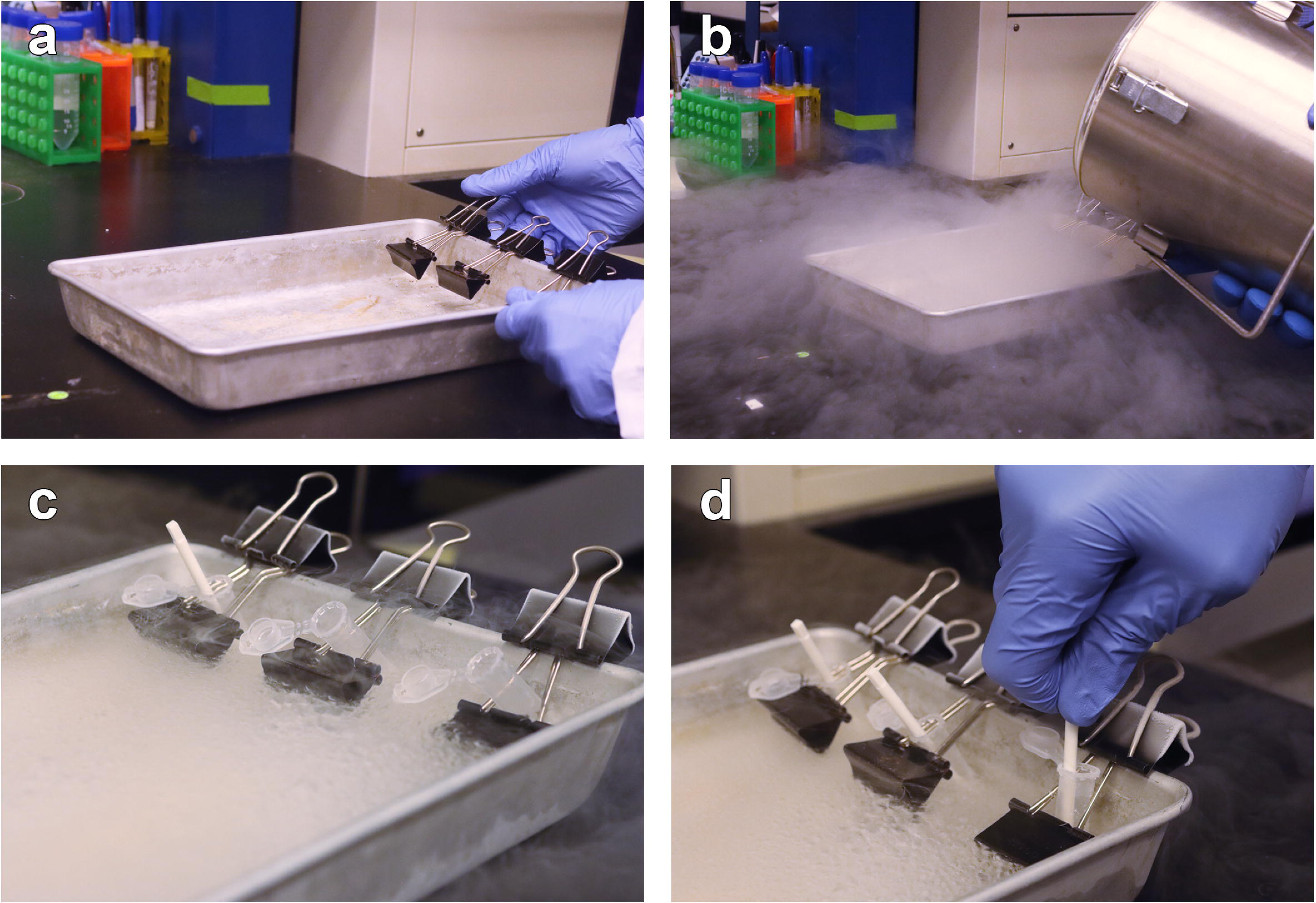

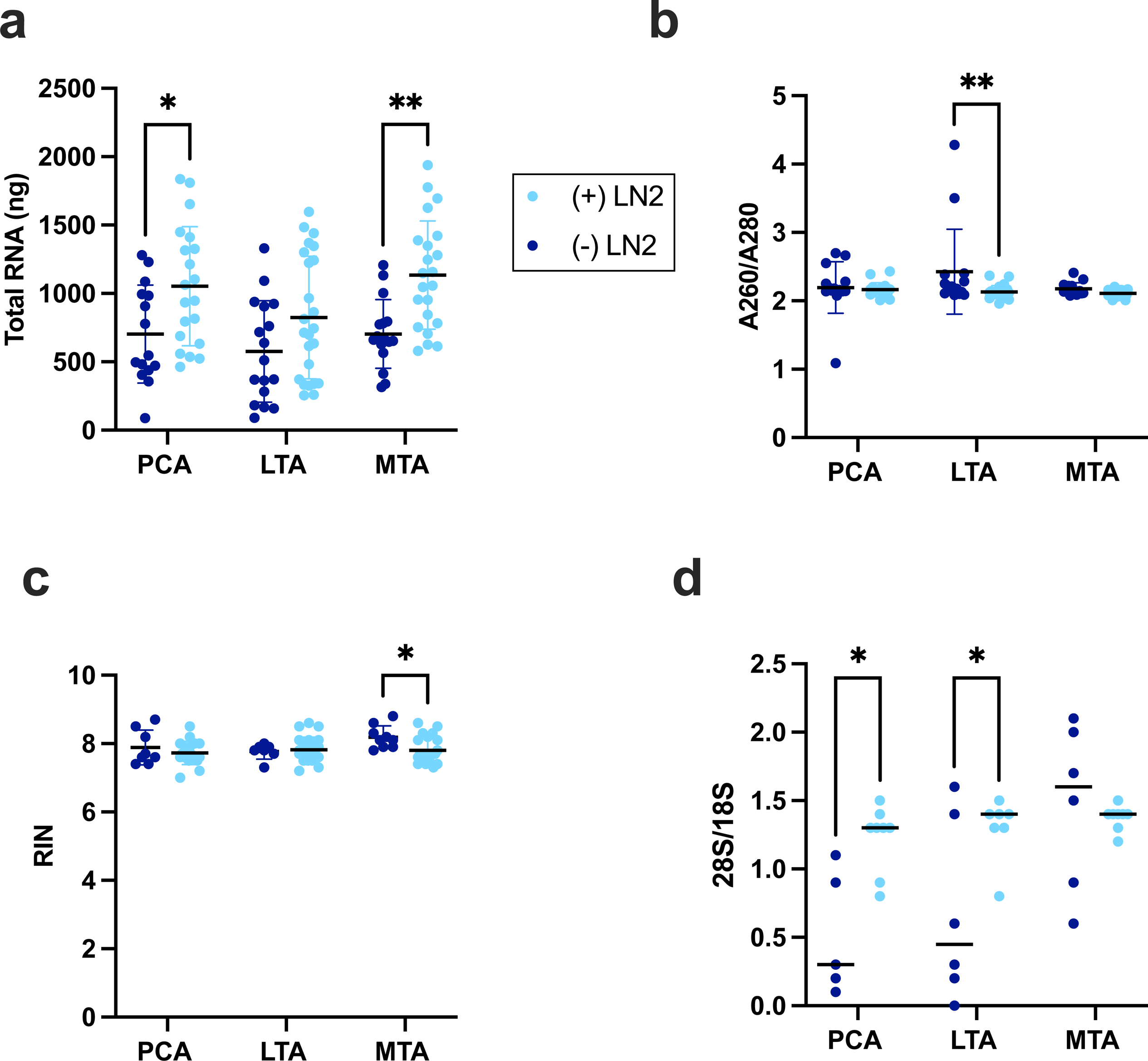

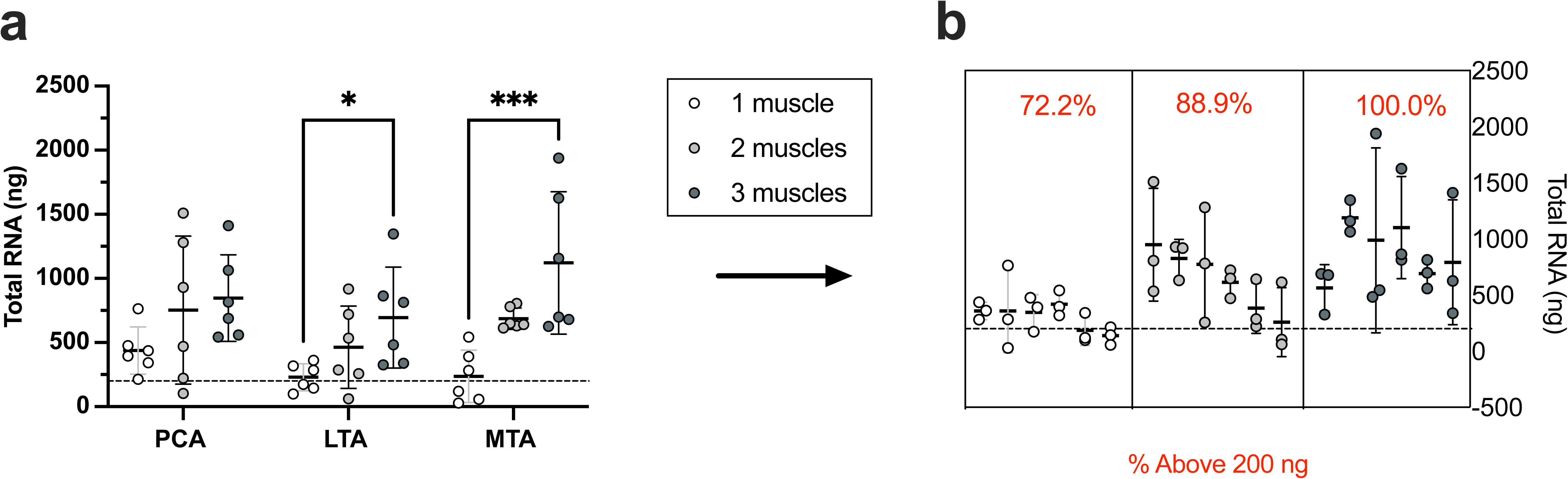

## Notes

### Competing Interest Statement

The authors have declared no competing interest.

