## Supplementary figures and images for "An optimized method for high-quality RNA extraction from distinctive intrinsic laryngeal muscles in the rat model"

### Supplementary Figure 1

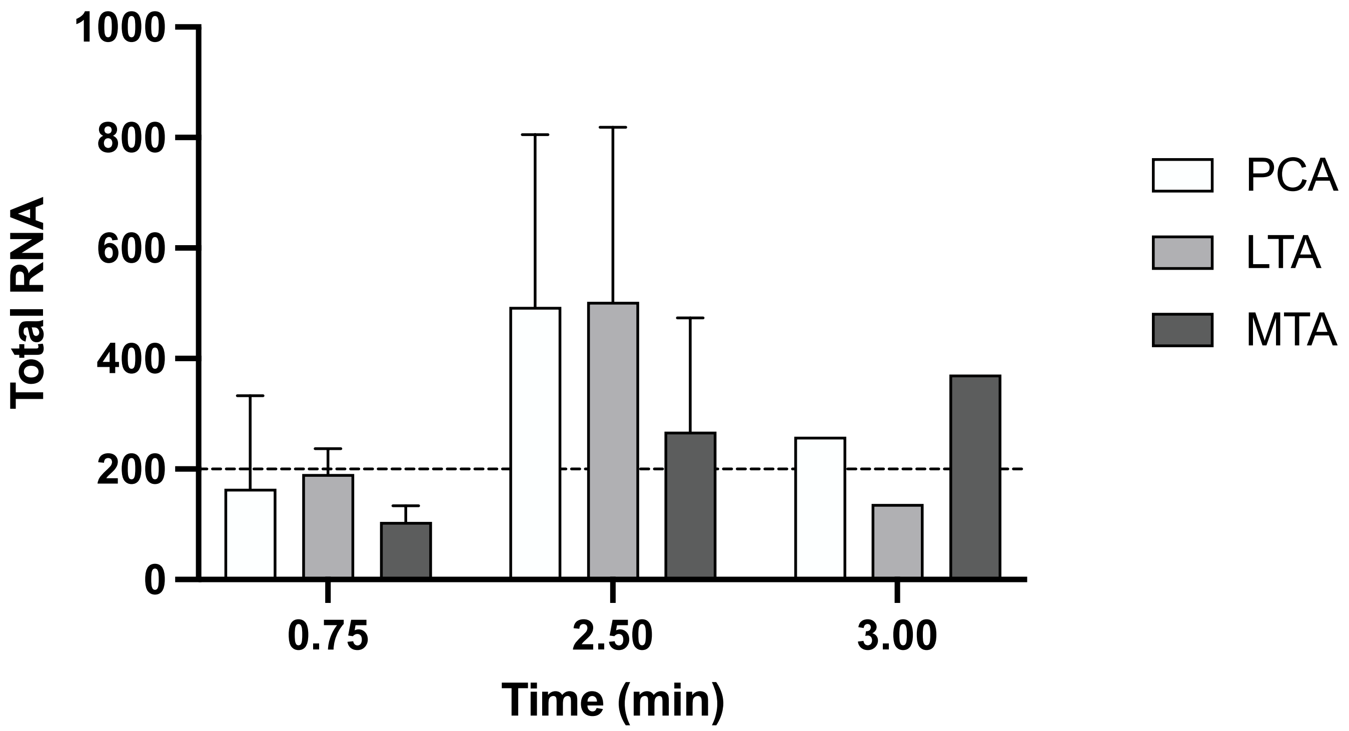

### Supplementary Figure 2

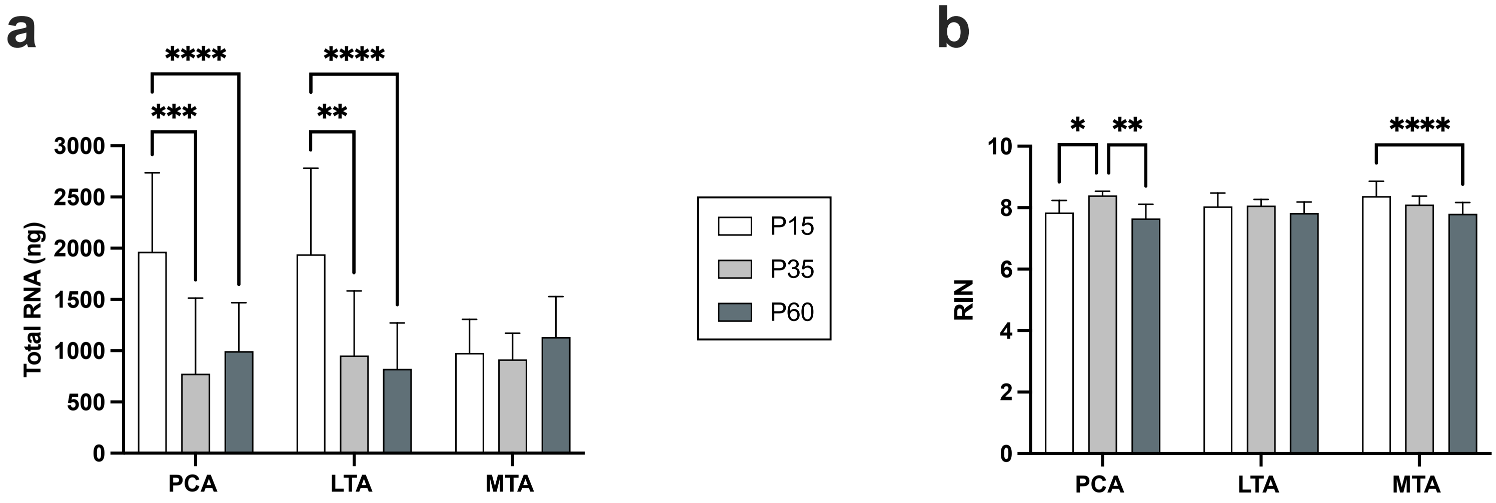
